# A near-complete genome assembly of the *Fusarium oxysporum* keratitis isolate MRL8996

**DOI:** 10.64898/2026.07.06.736765

**Authors:** Andrea Doddi, Gema Puebla Planas, Manuel Sánchez López-Berges, Antonio Di Pietro

## Abstract

*Fusarium oxysporum* MRL8996 is a fungal strain isolated from a severe case of contact lens-associated keratitis. Here we report a near-complete genome assembly of this isolate using a hybrid Nanopore and Hi-C scaffolding approach. The assembly resolves the genome into 16 distinct chromosomes, including 11 core and 5 lineage-specific chromosomes. This high-quality reference genome provides an unprecedented tool for investigating the large-scale structural variations and evolutionary mechanisms driving adaptation in this highly versatile fungal lineage.

## BACKGROUND & SUMMARY

The *Fusarium oxysporum* Species Complex (FOSC) encompasses a diverse group of soil-borne plant pathogens that also display a remarkable capacity for cross-kingdom pathogenicity^1,2^, causing opportunistic infections in mammals, including humans^3,4^. The clinical relevance of this pathogen is exacerbated by its broad-spectrum resistance to available antifungal agents^5,6^. Within the FOSC, strain MRL8996 (NRRL 47514), originally isolated from the contact lens of a patient with keratitis disease, represents an important clinical model^7^.

The evolutionary success and broad host range of *F. oxysporum* are generally attributed to its compartmentalised genome structure, comprising a combination of conserved core chromosomes and highly plastic accessory regions that have been associated with host specificity^8,9^. Population genomic studies indicate that a set of 11 core chromosomes is consistently conserved across the FOSC, whereas the accessory compartment exhibits significant structural variation and evolutionary dynamics across different isolates^2,8,10,11^. While the number and content of accessory chromosomes tend to be similar within a given plant-infecting *forma specialis*^12,13^, it has been suggested that in clinical strains such as MRL8996, accessory chromosomes could harbour genes involved in pre-adaptation to the human host^2^.

The currently available draft genome of MRL8996^2^(PRJNA554890) is fragmented into numerous contigs, complicating the study of large-scale genomic variations and host-adaptation mechanisms. Furthermore, while chromosome-level assemblies and Hi-C contact maps are now available for a few FOSC members^10,14–18^, high-resolution three-dimensional genomic data of clinical isolates have so far been lacking. Here we present a near-complete genome assembly of MRL8996, which resolves the complex nature of its accessory genome, providing a blueprint for investigating the genetic determinants that drive cross-kingdom adaptation.

The improved MRL8996 chromosome-level genome assembly was generated using a hybrid sequencing approach. Long-read Oxford Nanopore sequencing generated 4.56 Gb of high-quality reads, providing approximately 87× raw genome coverage (Table 1). Reads ≥ 10 kb were selected to achieve an optimum effective coverage for the initial *de novo* assembly, which was performed using Flye v. 2.9.6^19^. In addition, Hi-C scaffolding was incorporated to achieve chromosome-level continuity. The Hi-C library generated 20.39 Gb of raw data via 150-bp paired-end (PE150) sequencing (67,964,897 read pairs), providing an estimated 390× genome coverage. Sequencing coverage for both Nanopore and Hi-C reads was calculated based on the final assembled genome size of 52.32 Mb. The data were processed via Juicer v2.0^20^ to obtain 45,094,709 valid, unique chromatin interaction pairs (Table 1). Scaffolding was subsequently executed using 3D-DNA^21^ to construct the final pseudomolecules. As a final step, manual curation was carried out using Juicebox^22^.

**Table 1.** Assembly statistics and sequencing data of the *Fusarium oxysporum* MRL8996 genome.

| Assembly Features |  |
| --- | --- |
| Total assembly length (bp) | 52321866 |
| Total number of scaffolds | 16 |
| Resolved chromosomes (Core / Accessory) | 16 (11 / 5) |
| Largest scaffold (Mb) | 6.39 |
| Scaffold N50 (Mb) | 4.82 |
| Scaffold L50 | 5 |
| Overall GC content (%) | 47.66 |
| Sequencing Data |  |
| Oxford Nanopore reads (Gb) | 4.56 (~87× coverage) |
| Hi-C Illumina PE150 data (Gb) | 20.39 (~390× coverage) |
| Hi-C sequenced read pairs | 67964897 |
| Hi-C valid chromatin interactions | 45094709 |

The hybrid assembly pipeline produced 16 chromosome-scale scaffolds supported by high-depth Hi-C interaction blocks, totalling 52,321,866 bp. The use of Hi-C centromeric interaction blocks is standard in fungal genome analysis to define chromosome boundaries in cases where telomere-to-telomere assembly is not fully achieved^19^. In addition, StainedGlass^23^ was used to confirm the detection of centromeric regions. In total, 27 of the 32 telomeres were detected (Supplementary Image 1; tapestry v1.0.1^24^), representing a near-complete genome assembly. The assembly exhibits continuity metrics, with a scaffold N50 of 4.82 Mb, an L50 of 5, and the largest scaffold reaching 6.39 Mb (Figure 1A; Table 1; Supplementary Table 1). The overall genome architecture, visualised by circos plot, highlights a heterogeneous distribution of genomic features in line with a compartmentalised genome structure (Figure 1B), where the core chromosomes shared across the *F. oxysporum* species complex^8,11^ are clearly conserved. In contrast, the accessory chromosomes display a distinct genomic signature, characterised by a pronounced enrichment of transposable elements (TEs) and increased simple repeat density; a substantial decrease in overall gene and exon densities; and a complete absence of biosynthetic gene clusters (BGCs), which are widely distributed across the core genome.

**Figure 1.**
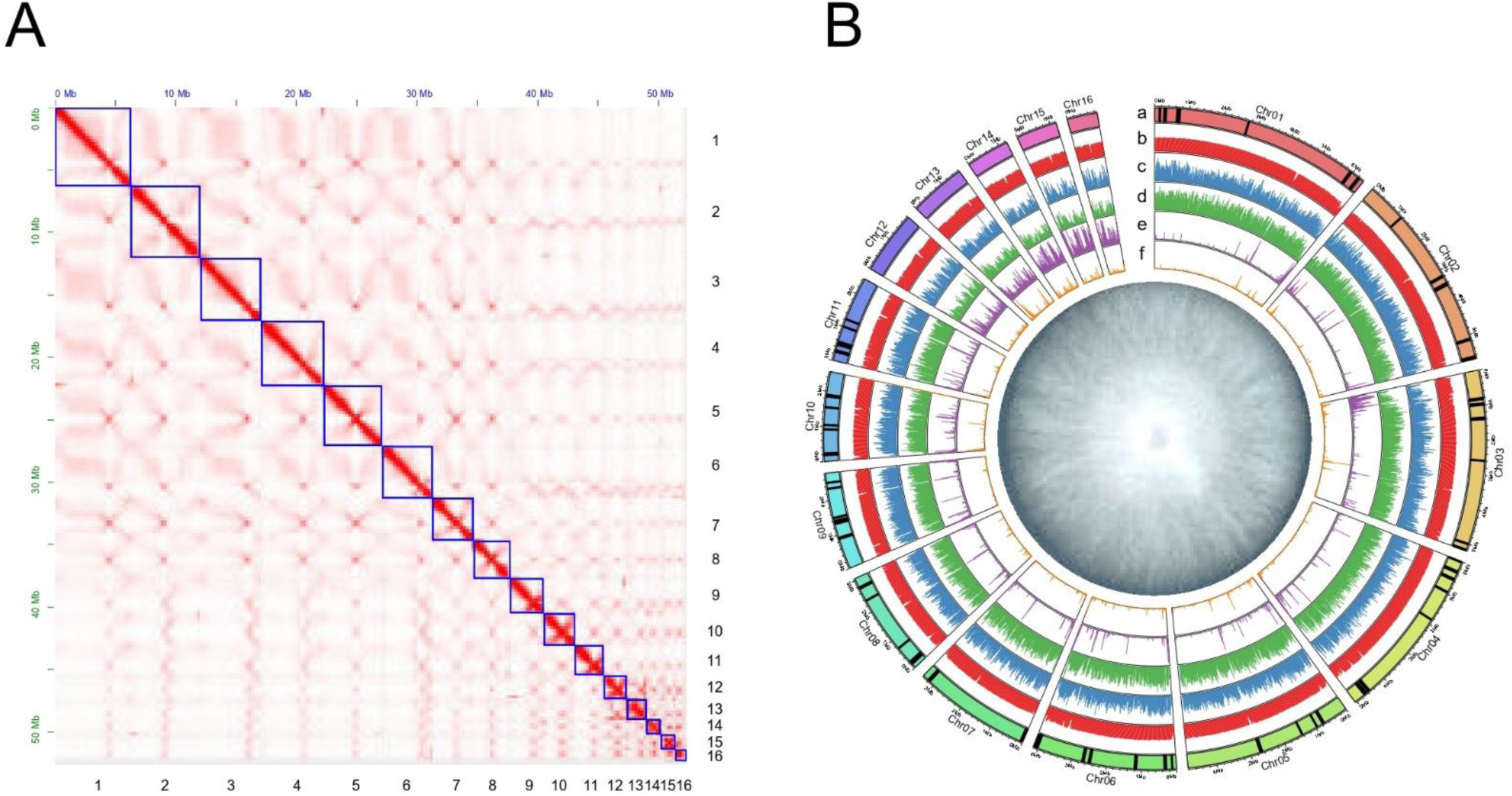
Overview of the chromosome-level genome assembly and annotation of *Fusarium oxysporum* strain MRL8996. **(A)** Genome-wide Hi-C contact map showing the interaction matrix among the 16 assembled chromosomes. **(B)** Circos plot illustrating the genomic features of the MRL8996 genome. Tracks from outer to inner represent: (a) Ideograms of the 16 chromosomes, in which the black vertical lines represent the genomic locations of biosynthetic gene clusters (BGCs); (b) GC content; (c) gene density; (d) exon density; (e) repetitive DNA (transposable elements) density; and (f) simple repeat density. Genomic feature densities (tracks b to f) were calculated in non-overlapping 20 kb windows and represented on a scale from 0 to 1, except for track f, which ranges from 0 to 0.2. At the centre, a picture of the MRL8996 strain growing in a PDA plate at 5 days post-inoculation. Chr: chromosome

A direct comparison with the previously available draft genome sequence reveals a significant improvement in contiguity for the present assembly (Figure 2; Table 2). While the earlier MRL8996 v1.0 assembly from the JGI genome portal database^2^ (PRJNA554890) consisted of 250 contigs with a scaffold N50 of 1.73 Mb, an N90 of 0.13 Mb, and L50/L90 values of 10 and 46, respectively (Table 2), the Hi-C-based chromosome-level assembly reported here (v2.0) condenses the genome into only 16 pseudomolecules, representing a significant reduction in the number of contigs. The new assembly exhibits a scaffold N50 of 4.82 Mb, 2.8 times higher than that of v1.0, along with an improved N90 of 1.82 Mb. The L50 and L90 values decrease from 10 to 5 and from 46 to 12, respectively (Table 2). Additionally, the largest assembled sequence increases from 4.65 Mb to 6.39 Mb (Table 2).

**Figure 2.**
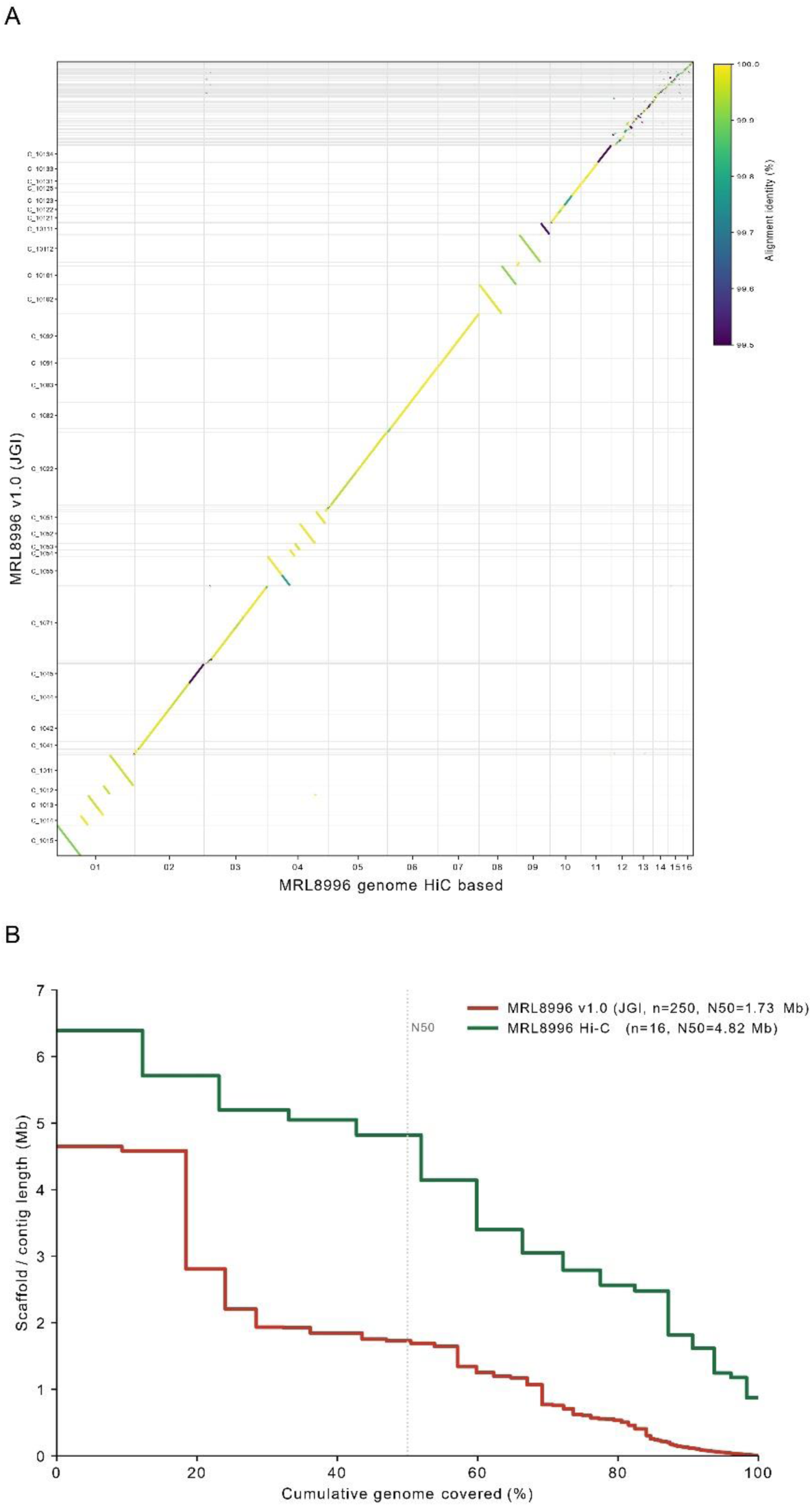
Comparison of genome assemblies v1.0 and v2.0 of *Fusarium oxysporum* MRL8996. (A) D-GENIES dotplot alignment comparing the chromosome features between the highly fragmented JGI v1.0 draft assembly (y-axis) and the newly generated Hi-C-based chromosome-level assembly (x-axis). The colour scale indicates the percentage of alignment identity. (B) Cumulative length plot comparing the structural contiguity of the two assemblies. The graph illustrates scaffold and contig lengths in megabases (Mb) against the cumulative percentage of the genome covered. The red line represents the baseline MRL8996 v1.0 assembly (n=250, N50=1.73 Mb), while the green line indicates the Hi-C scaffolding (n=16, N50=4.82 Mb). The vertical dashed line indicates the N50 metric threshold.

**Table 2.** Comparison of assembly statistics for the two genome versions of *Fusarium oxysporum* MRL8996.

| **Statistic** | **MRL8996 v1.0**<br>**(JGI)** | **MRL8996 v2.0**<br>**(this study)** | **Difference percentage**<br>**(%)** |
| --- | --- | --- | --- |
| Number_of_contigs | 250 | 16 | -93.6 |
| Assembly_size_bp | 50071586 | 52321866 | 4.5 |
| N50_bp | 1727337 | 4819956 | 179 |
| N90_bp | 129585 | 1818122 | 1303 |
| L50 | 10 | 5 | -50 |
| L90 | 46 | 12 | -73.9 |
| Largest_contig_bp | 4647901 | 6391895 | 37.5 |
| GC_percent | 47.6 | 47.66 | 0.1 |
| Ns_per_100kbp | 0.04 | 0.57 | NA |

Whole-genome alignment of the two assemblies using D-GENIES (Figure 2A) demonstrated that the 250 contigs from the v1.0 assembly align with the 16 chromosome-scale scaffolds of v2.0 with near-identical sequence identity. The main improvement of the v2.0 assembly is the high resolution of the accessory chromosomes, where most of the small contigs from the v1.0 assembly are positioned (Figure 2A). The assembly consolidates the previously scattered contigs into contiguous chromosomes without apparent large-scale rearrangements or loss of sequence content. The consistent base composition of the two versions further supports the integrity of the new assembly (Table 2).

Total assembly length between v1.0 and v2.0 increased by approximately 2.25 Mb, from 50.07 Mb to 52.32 Mb, likely reflecting an improved resolution and placement of repetitive and accessory sequences that were collapsed or remained unassembled in the previous draft. The small rise in ambiguous bases (0.04 to 0.57 Ns per 100 kbp) corresponds to the gap-spanning N runs introduced between adjacent contigs during Hi-C scaffolding (Supplementary Table 1). Together, these metrics establish the v2.0 genome assembly as a substantially more contiguous and complete reference for the study of large-scale structural variations and accessory chromosome content.

To further resolve the compartmentalised architecture of the MRL8996 genome, we compared its macrosynteny against the tomato pathogenic reference strain *F. oxysporum* f. sp. *lycopersici* 4287 (GCA_000149955.2^25^) and the endophytic reference strain Fo47 (GCA_013085055.1^26^) (Figure 3A). In addition, we examined internal sequence redundancy through self-alignment using minimap2^27^ (Figure 3B). We found that the first 11 pseudomolecules, corresponding to Chr01–Chr11, aligned one-to-one with the core chromosomes of Fol4287 and Fo47, predominantly in the same order and covering most of their lengths. This clearly identified Chr01–Chr11 as the core chromosome complement of the MRL8996 genome, which is conserved across the FOSC^8,9,11^. In contrast, the remaining five MRL8996 chromosomes (Chr12–Chr16) failed to align with a single orthologous counterpart. Instead, they exhibited small partial alignment matches, confined primarily to the lineage-specific Fol4287 chromosomes 03, 06, 14 and 15, and to the small accessory Chr07 of Fo47, but not to the conserved core genome (Figure 3A, Supplementary Tables 2 and 3). Self-alignment further reinforced this distinction: the core chromosomes appeared essentially as a single copy with minimal inter-chromosomal similarity, whereas the accessory chromosomes shared mutual sequence homologies (Figure 3B). Shared homologies were most pronounced among MRL8996 Chr14-Chr16, including repeat-derived segments that are also present in the core, indicating a mosaic duplication- and repeat-rich organisation (Supplementary Table 4).

**Figure 3.**
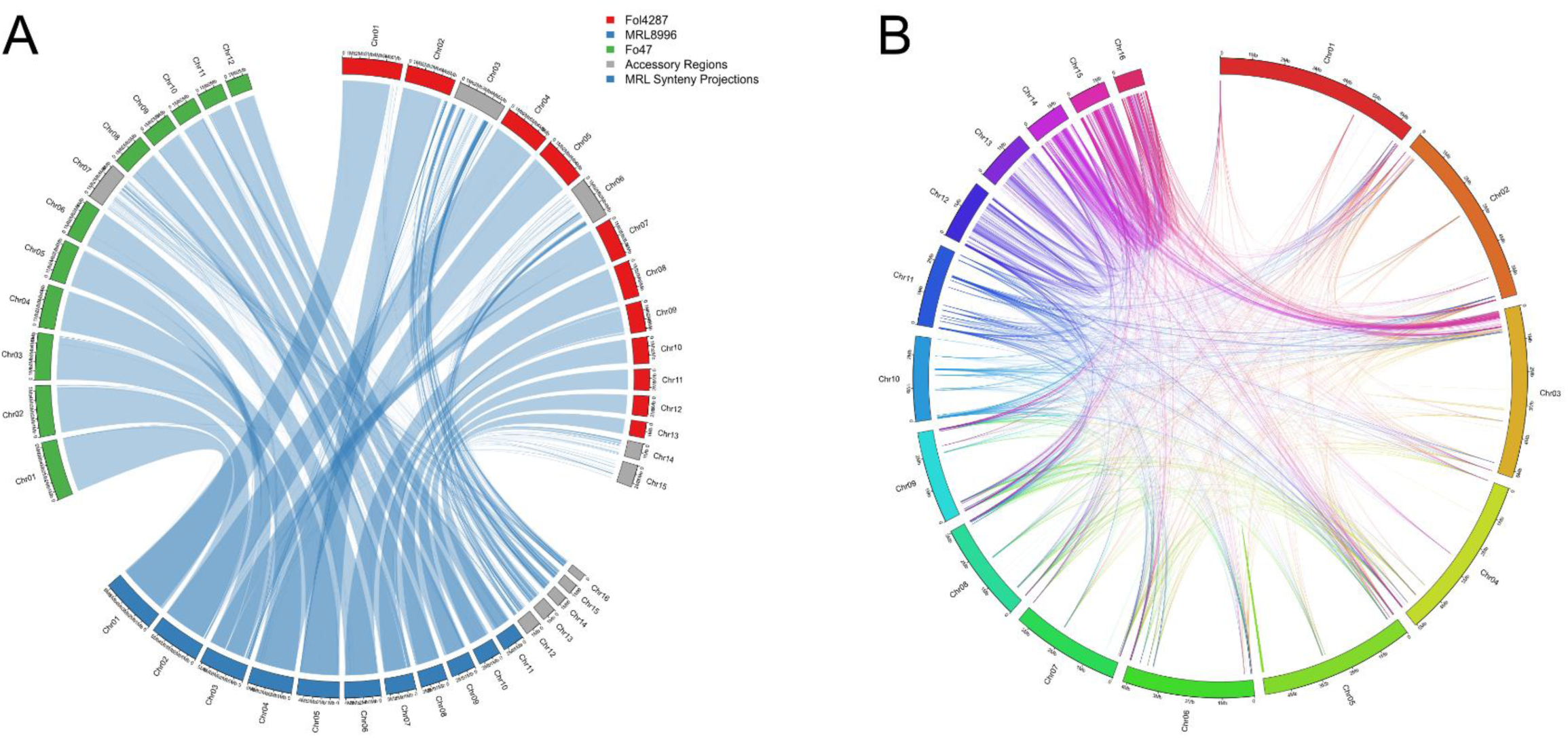
Inter- and intra-genomic synteny of *Fusarium oxysporum* MRL8996. **(A)** Circos plot comparing the genome architectures of *F. oxysporum* MRL8996 (bottom, blue), the tomato pathogenic reference strain Fol4287 (right, red), and the endophytic biocontrol strain Fo47 (left, green). Bars represent individual chromosomes, while blue ribbons represent syntenic blocks based on the MRL8996 genome. **(B)** The circos plot illustrates intra-genomic duplications within the MRL8996 genome. The outer ring displays the MRL8996 chromosomes, while the internal colored links represent extensive self-alignments and segmental duplications.

**Table 3.** Statistics of repeat sequences in the *Fusarium oxysporum* MRL8996 genome.

| TE_Class | Total bp | Percentage of the genome (%) |
| --- | --- | --- |
| unknown | 638833 | 1.221 |
| LTR/Gypsy/Ty3 | 554888 | 1.0605 |
| ClassI_LTR_Gypsy | 424182 | 0.8107 |
| LINE/Tad1 | 370859 | 0.7088 |
| LTR/Gypsy | 333285 | 0.637 |
| DNA/Helitron | 328240 | 0.6273 |
| DNA/DTM | 237076 | 0.4531 |
| ClassII_DNA_TcMar_nMITE | 204686 | 0.3912 |
| LTR/Copia | 166699 | 0.3186 |
| DNA/DTC | 161420 | 0.3085 |
| SDR/Crypton | 160890 | 0.3075 |
| LTR/unknown | 156603 | 0.2993 |
| ClassIII_Helitron | 139981 | 0.2675 |
| TIR/Tc1/mariner | 132154 | 0.2526 |
| TIR/hAT | 120973 | 0.2312 |
| ClassII_DNA_CACTA_nMITE | 110611 | 0.2114 |
| ClassII_DNA_Harbin-ger_nMITE | 105872 | 0.2023 |
| DNA/DTH | 100154 | 0.1914 |
| Helitron/Helitron | 78460 | 0.15 |
| ClassII_nMITE | 72329 | 0.1382 |
| ClassI | 64483 | 0.1232 |
| ClassI_nLTR_DIRS | 64272 | 0.1228 |
| ClassII_MITE | 63467 | 0.1213 |
| ClassII_DNA_TcMar_MITE | 62036 | 0.1186 |
| LINE/CRE | 55285 | 0.1057 |
| ClassI_LTR_Copia | 41357 | 0.079 |
| ClassI_nLTR_SINE_tRNA | 35857 | 0.0685 |
| ClassI_nLTR_LINE_L1 | 35321 | 0.0675 |
| SINE/Foxy | 30920 | 0.0591 |
| DNA/DTT | 26457 | 0.0506 |
| TIR/Mule | 24878 | 0.0475 |
| ClassII_DNA_Harbin-ger_MITE | 24852 | 0.0475 |
| Unknown/Unknown | 22785 | 0.0435 |
| ClassII_DNA_hAT_nMITE | 19436 | 0.0371 |
| ClassII_DNA_hAT_MITE | 19288 | 0.0369 |
| DNA/PiggyBac | 17573 | 0.0336 |
| ClassII_DNA_CACTA_MITE | 16383 | 0.0313 |
| DNA/DTA | 15200 | 0.0291 |
| DNA/hAT-Restless | 11903 | 0.0227 |
| ClassI_LTR | 11185 | 0.0214 |
| Tac/Tac | 8740 | 0.0167 |
| MITE/DTM | 7488 | 0.0143 |
| Marsu/Marsu | 6666 | 0.0127 |
| ClassI_nLTR | 6633 | 0.0127 |
| ClassII_DNA_Mutator_nMITE | 5931 | 0.0113 |
| ClassII_DNA_Mutator_MITE | 2076 | 0.004 |
| SINE? | 1855 | 0.0035 |
| MITE/DTC | 1570 | 0.003 |
| ClassII_DNA_TcMar_unknown | 1089 | 0.0021 |
| tRNA | 846 | 0.0016 |
| MITE/DTA | 737 | 0.0014 |
| ClassII_DNA_hAT_unknown | 564 | 0.0011 |
| MITE/DTH | 505 | 0.001 |
| ClassI_nLTR_SINE | 284 | 0.0005 |
| LINE/Gypsy/Ty3 | 267 | 0.0005 |
| MITE/DTT | 118 | 0.0002 |
| ClassI_nLTR_SINE_7SL | 108 | 0.0002 |
| Total | 5306610 | 10.1418 |

Overall, these findings define 11 stable core chromosomes and 5 plastic accessory chromosomes, representing the conserved and variable components, respectively, of the MRL8996 genome. This provides a structural framework for investigating the accessory-encoded virulence determinants on the human host.

Beyond chromosome-level architecture, the MRL8996 genome encodes a large repertoire of predicted secreted proteins that may be relevant for its lifestyle. Using the JGI annotation of the protein-filtered model, we predicted secreted proteins independently with SignalP^28^ and EffectorP^29^. The matching results were then checked for the absence of a transmembrane domain using DeepTMHMM^30^, ultimately compiling a catalogue of 513 secreted candidates widely distributed across the genome with a lower prevalence in accessory chromosomes (Supplementary Image 2). Since effector candidates often lack detectable sequence homology while converging on a limited set of protein folds, we classified the effectors based on their structural characteristics rather than their sequence^31,32^

Three-dimensional models for the 513 candidate effectors were predicted using ColabFold AlphaFold2 v.1.6.1^33^, and the resulting structures were compared in an all-versus-all manner. Of the 513 candidates, 482 were selected based on a threshold of pLDDT >50, while the remaining 31 structures were excluded. Under a TM-score cutoff of 0.5, hierarchical clustering (Figure 4A) resolved the 482 predicted effectors into 22 structural families (≥5 members), consistent with the independent DALI network organisation (Figure 4B). This grouping was established by cutting the dendrogram to a height of 0.5 (i.e. a TM-score similarity threshold >0.5), which approximates fold-level similarity. The exact number of structural families depends on the TM-score cutoff and linkage; therefore, the reported count should be interpreted within this analytical framework. The five largest families comprised 26, 21, 18, 14 and 13 members, respectively (Supplementary Table 5).

**Figure 4.**
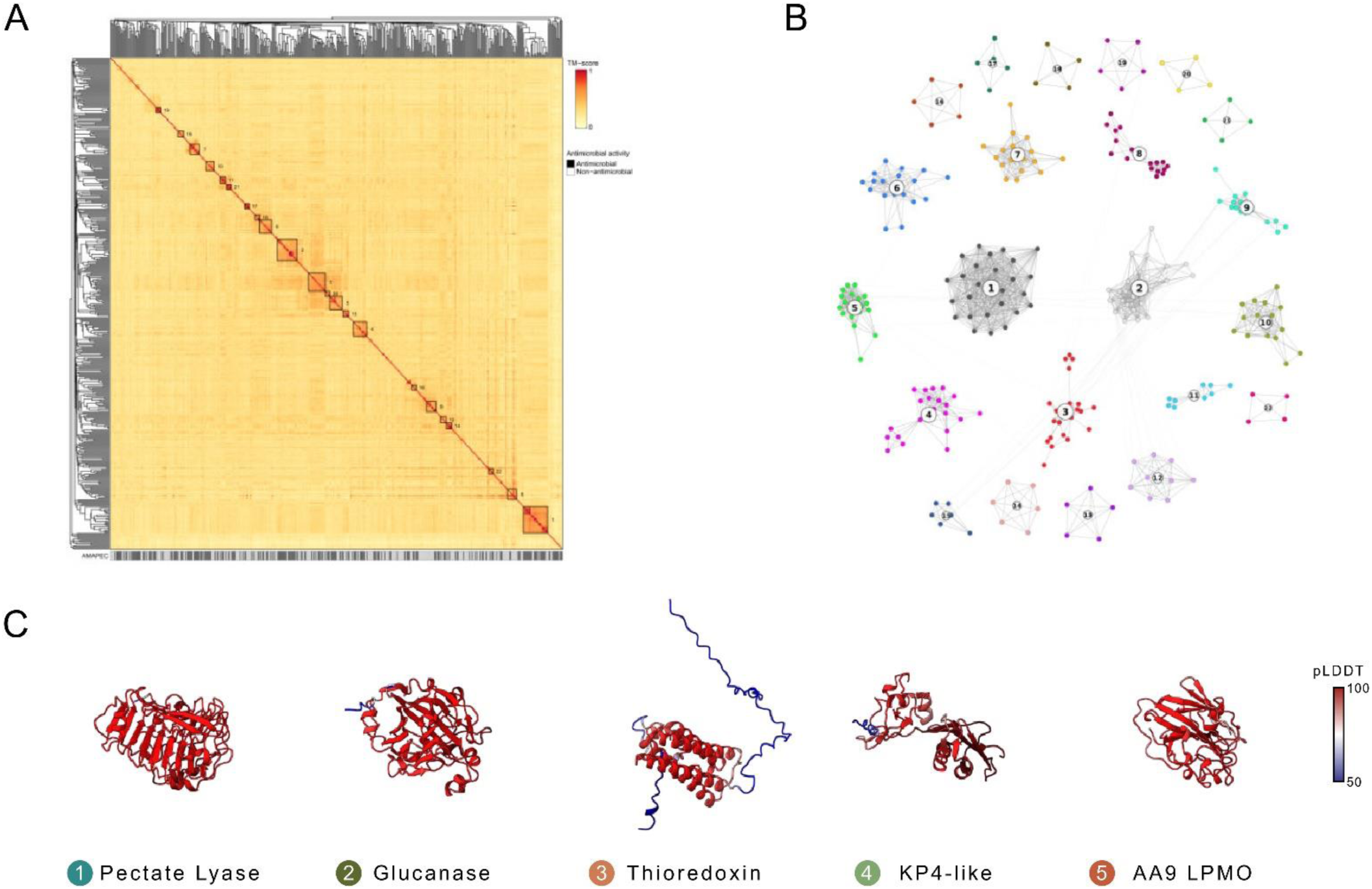
Structure-based classification of the predicted effector repertoire of *Fusarium oxysporum* MRL8996. **(A)** Heatmap of all-versus-all pairwise structural similarity (TM-score; colour scale 0–1) for the 482 effector candidates with a predicted-model confidence pLDDT > 50, ordered by hierarchical clustering (average linkage / UPGMA of the 1 − TM-score distance matrix). The 22 structural families with at least 5 members are outlined along the diagonal and numbered in order of decreasing size. The annotation track beneath the heatmap shows the per-effector AMAPEC antimicrobial prediction (black, predicted antimicrobial; grey/white, non-antimicrobial). **(B)** Force-directed structural-similarity network of the same 482 effectors. Nodes represent effectors and edges connect structurally similar pairs with a DALI Z-score ≥ 5.2, nodes with a degree < 3 were excluded. Communities detected with the Louvain algorithm are shown in distinct colours and numbered according to decreasing size. Note that the Louvain communities in (B) were partitioned independently from the hierarchical families in (A), although the largest groups broadly correspond. The graph was laid out with the Kamada–Kawai algorithm. **(C)** Representative ColabFold (AlphaFold2) models of the five largest structural families defined in panel A, selected as the structural medoid of each family, colored by per-residue confidence (pLDDT; scale 50–100): (1) pectate lyase, (2) glucanase, (3) thioredoxin, (4) KP4-like and (5) AA9 lytic polysaccha-ride monooxygenase (LPMO).

To further validate these groupings, an independent force-directed structural-similarity network (Figure 4B) was constructed, revealing a similar broad organisation that partitioned the effectors into discrete, densely connected communities separated by few inter-community links (Supplementary Table 6). The five most populated families correspond to well-defined fungal effector folds: pectate lyase^34^ (Family 1), glucanase^35^ (Family 2), thioredoxin^36^ (Family 3), KP4-like^37^ (Family 4) and AA9 lytic polysaccharide monooxygenase^38^ (LPMO; Family 5) (Figure 4C).

Growing evidence suggests that fungal effectors may have originated as antimicrobial proteins^39–41^ that are deployed to manipulate competing microbes, as reported in several soil-borne fungi such as *V. dahliae*^42–46^ and *Rosellinia necatrix*^47^. To identify candidates with potential roles in microbial antagonism, the 482 structurally characterised effectors were scored for predicted antimicrobial activity using AMAPEC^39^ and annotated on the heatmap (Figure 4A), revealing 241 potential antimicrobials that were particularly enriched in the pectate lyase and glucanase families. Together, these analyses provide a structural and functional map of the putative MRL8996 effector proteins that complement the sequence-based annotations and support further investigation into the determinants of cross-kingdom interactions.

In summary, this study presents an improved, near-complete genome assembly of the clinical *F. oxysporum* isolate MRL8996, offering a high-resolution view of its structural organisation within this lineage. These findings advance our understanding of genome compartmentalisation and evolution in the FOSC, establishing a solid foundation for exploring the molecular mechanisms underlying human infections by *F. oxysporum*.

## MATERIALS AND METHODS

### Fungal cultures, DNA extraction, Nanopore library preparation and sequencing

The MRL8996 strain used in this study was originally isolated from a patient with contact lens-associated keratitis (Cleveland Clinic Foundation, Ohio, USA, 2006). An aliquot of frozen microconidia taken from the glycerol stock was grown in 100 ml of homemade potato dextrose broth (PDB) culture in a 500 ml flask for 3 days at 28 °C. For genomic DNA isolation, mycelium was collected by filtration through a Monodur nylon filter and freeze-dried overnight. High-molecular-weight (HMW) DNA was extracted following the protocol of Chavarro-Carrero et al. (2021). DNA quality, size, and quantity were assessed using Nanodrop, Qubit, and gel electrophoresis. Library preparation with the Ligation Sequencing Kit (SQK-LSK114) was performed with ∼1.5 μg HMW DNA according to the manufacturer’s instructions (Oxford Nanopore Technologies, Oxford, UK). An R10.4.1 flow cell (Oxford Nanopore Technologies, Oxford, UK) was loaded and run for 24 h, and subsequent base calling was performed using Dorado (version 7.11.2; Oxford Nanopore Technologies, Oxford, UK). To obtain longer reads, ultra-high-molecular-weight (UHMW) DNA, an extra clean-up step was performed. To this end, after DNA precipitation, the Short Reads Eliminator Kit (PacBio, California, USA) was used following the manufacturer’s protocol to select DNA fragments >25 kb.

For Hi-C sequencing, an aliquot of MRL8996 spores was inoculated into 400 ml of PDB in a 1500 ml flask for 3 days at 28 °C. The resulting biomass (mycelium and spores) was harvested by centrifugation at 10000 × g for 10 minutes, followed by two consecutive washes with sterile distilled water. Three grams of the sample were ground to a fine powder using liquid nitrogen. The pulverised tissue was resuspended in 10 ml PGShield, provided by Phase Genomics (Seattle, WA, USA), to ensure sample stabilisation for ambient-temperature shipping.

### *De novo* genome assembly and chromosome-level assembly

To obtain a first draft assembly, the reads from Nanopore sequencing were used to generate a *de novo* assembly of MRL8996 using Flye v.2.9^19^ with -nano-hq -g 50m -m 10000 parameters to verify the assembly quality and determine the final genomic sequence. Redundant contigs were deleted, and the corrected assembly was generated as output. To improve the draft assembly, Hi-C sequencing data were generated to verify and produce a chromosome-level genome assembly. Hi-C library construction of *F. oxysporum* strain MRL8996 was performed according to the standard protocol and sequenced on the Illumina NovaSeq 6000 platform (Phase Genomics Inc., Seattle, WA, USA). Hi-C reads were mapped onto this assembly with Juicer (v2.0)^20^ using the “assembly” option to skip the post-processing steps and generate the merged_nodups.txt file. For the Juicer pipeline, restriction site maps were generated using the DpnII (GATC) restriction site profile and the assembly was indexed with BWA index (v0.7.17-r1188)^49^ and used to polish the assembly using 3D-DNA^21^. Subsequently, Juicebox (v1.11.08)^22^ was used to manually curate the genome assembly. Contigs were merged into scaffolds according to the Hi-C map, and Ns were introduced between contigs within scaffolds; gaps between contigs were removed, and contigs were merged. Ultimately, TGS-GapCloser v1.2.1^50^ was used to close the remaining gaps, resulting in 1 gap closure and 3 gaps remaining.

### Gene annotation and repetitive element analysis

Gene models for the chromosome-level assembly were obtained from the JGI genome portal deposition^51^, lifted over from the MRL8996 v1.0 draft using liftoff v1.6.3^52^, and only the filtered ‘best’ gene models were retained for downstream analyses. First, repetitive sequences and transposable elements (TEs) were *de novo* predicted and classified using the Extensive *de-novo* TE Annotator (EDTA) pipeline v2.2.2^53^. To enhance the accuracy of TE identification, the *de novo* predictions were supplemented with a thoroughly curated TE library derived from the reference strain *F. oxysporum* f. sp. *lycopersici* 4287 (Fol4287)^8^. The resulting consolidated, non-redundant TE library was subsequently used to soft mask the MRL8996 genome assembly using the RepeatMasker^54^ engine integrated into the EDTA framework (Table 3, Supplementary Table 7). Ultimately, to better characterise TEs predicted by the EDTA pipeline, DeepTE^55^ was used to categorise unknown elements from EDTA prediction. In addition, Kimura divergence was calculated to predict active TEs (Supplementary Image 3). Biosynthetic gene clusters (BGCs) were predicted with antiSMASH fungal v8.0^56^. For the genome overview shown in Figure 1B, GC content, gene density, exon density, transposable-element density and simple-repeat density were computed in non-overlapping 20-kb windows and plotted together with the BGC positions using the circlize R package v0.4.18^57^.

### Effector prediction and structural classification

To identify the effector repertoire of MRL8996, we established a predictive pipeline similar to that recently developed for the gap-free genome assembly of the *F. oxysporum* f. sp. *conglutinans* strain Fo5176^16^ and also used for the *F. oxysporum* biocontrol strain FO12^17^. Secreted protein candidates were predicted from the filtered best gene models of the JGI annotation. We used EffectorP v3.0^29^ and SignalP v.6.0^28^ to independently identify potential secreted proteins within the predicted proteome of MRL8996, and we compiled the effectors that were common to both predictions (Supplementary Table 8-9). The candidate effectorome was subsequently analysed using deepTMHMM v.1.0^30^ to filter out the candidates with transmembrane helices, resulting in the final set of predicted secreted effectors (Supplementary Table 10).

Three-dimensional models of all 513 effector candidates were predicted using ColabFold v1.6.1 (AlphaFold2)^33^, with default settings except for –models 3. Per-residue model confidence (pLDDT) was retained for downstream visualisation. Pairwise structural similarity was computed all-versus-all with TM-align v0.3.0^58^. For each pair, the higher of the two length-normalised TM-scores was retained (Max_TM_Score, Supplementary Table 11), giving a symmetric 482 × 482 similarity matrix for the candidates with successful comparisons (31 of the 513 candidates were not represented in the matrix because of their low pLDDT score < 50). For the structural heatmap (Figure 4A), the TM-score matrix was converted to a distance matrix (1 − TM-score) and clustered hierarchically using average linkage^59^ (UPGMA). Structural families were defined by cutting the dendrogram at a height of 0.5 (i.e. a TM-score similarity threshold > 0.5; cutree, RStudio (v4.6.1). Families with at least 5 members were outlined and numbered in order of decreasing size. The heatmap was rendered with the ComplexHeatmap package v2.28.0^60^ in RStudio.

A structural-similarity network (Figure 4B) was constructed independently of the heatmap. Nodes represent effectors and edges connect pairs with a DALI Z-score ≥ 5.2, derived from the empirical Z-score distribution rather than a literature value. To filter out background noise, nodes with a degree <3 were removed^31^. Communities were detected with the Louvain algorithm, and the graph was laid out with the Kamada–Kawai algorithm; network analysis and visualisation were performed with the igraph v2.3.2^61^ and ggraph v2.2.2^62^ packages in RStudio. Representative models for the five largest effector families relative to the structural network (Figure 4C) were selected as the structural medoid of each family (defined as the member with the highest summed within-family Z-score) and rendered in ChimeraX v1.12^63^, coloured by per-residue pLDDT (scale 50–100). Family identities (pectate lyase, glucanase, thioredoxin, KP4-like, AA9 LPMO) were assigned by Foldseek^64^ search against PDB, InterPro/Pfam of the family members, and structural inspection.

Antimicrobial activity was predicted with AMAPEC v1.0^39^ (https://github.com/fantin-mesny/amapec), which classifies effector candidates according to sequence, physicochemical properties, and predicted three-dimensional structure; the same ColabFold AlphaFold2^33^ models were supplied as input. The binary prediction (antimicrobial / non-antimicrobial) was displayed as an annotation track beneath the heatmap (Figure 4A, Supplementary Table 12).

### Data Records

The raw sequencing data of Nanopore and HiC have been deposited in the National Centre for Biotechnology Information (NCBI) under the BioProject number PRJNA554890 with the accession numbers SRR39391754^65^ SRR39391753^66^, respectively. The final assembled genome is deposited under the same BioProject at NCBI (GCA_009746015.2).

### Technical Validation

#### Manual correction, validation and evaluation of the genome assembly

The obtained genome assembly was manually corrected by removing redundant contigs and correcting the mis-assemblies using Hi-C read alignment within Juicebox visualisation^22^. Foreign Contamination Screening (FCS) NCBI tool was run to ensure the absence of contaminant sequences, such as the mitochondrial genome and sequencing adaptors. We utilised StainedGlass^23^ and tapestry v1.0.1^24^ to position all centromeres and almost all telomeres (27/32) (TTAGGG) (Supplementary Image 1).

#### Assessment of assembly quality and completeness

The accuracy and completeness of the assembly were evaluated by performing a BUSCO analysis (v.6.1.0)^67^ against the Hypocreales database. This reported a result of 99.6%, confirming the high quality of the genome data. To assess the base-level precision and structural fidelity of the assembly, a reference-free k-mer spectra analysis was executed using Merqury v.1.3^68^. The genome of *F. oxysporum* MRL8996 was compared against a high-accuracy k-mer database (k=21) generated from the Nanopore sequencing reads. Through this approach, the assembly achieved a consensus Quality Value (QV) of 54.10, while the total k-mer completeness was estimated to be 99.46% (Supplementary Image 4).

## Data availability

The genome assembly of *F. oxysporum* strain MRL8996 has been deposited in NCBI GenBank under the identifier GCA_009746015.2. The Nanopore and Hi-C sequencing data that support this genome assembly are available in the Sequence Read Archive (SRA) with the run numbers SRR39391754 ([Nanopore sequencing data - SRA - NCBI](https://www.ncbi.nlm.nih.gov/sra/SRX34109262%5baccn%5d)) and SRR39391753 ([HiC sequencing data - SRA - NCBI](https://www.ncbi.nlm.nih.gov/sra/SRX34109263%5baccn%5d)). Both datasets are associated with the BioProject number PRJNA554890 (https://www.ncbi.nlm.nih.gov/bioproject/PRJNA554890).

## Code availability

All commands and pipelines used in data processing were executed according to the manual and protocols of the corresponding bioinformatic software, and described in the Methods section, along with the versions. If no detailed parameters were mentioned for the software in this study, default parameters were used as suggested by the developer. All command-line strings, software parameters, and parsing pipelines used for the *de novo* genome assembly, scaffolding, structural/functional annotation, and effector prediction are publicly available in the GitHub repository : https://github.com/doddipetto/MRL8996_genome_assembly.

## AUTHOR CONTRIBUTIONS

AD and GPP conceived the project. AD designed the experiments. Phase Genomics Inc. performed Hi-C library and sequencing. AD and GPP analysed the data. ADP and MSL-B provided funding. AD and GPP wrote the manuscript. MSL-B and ADP revised the manuscript. All authors read and approved the final manuscript.

## ACKNOWLEDGMENTS

We thanks Prof. Li-Jun Ma for providing *Fusarium oxysporum* isolate MRL8996

---

**Supplementary Image 1.**
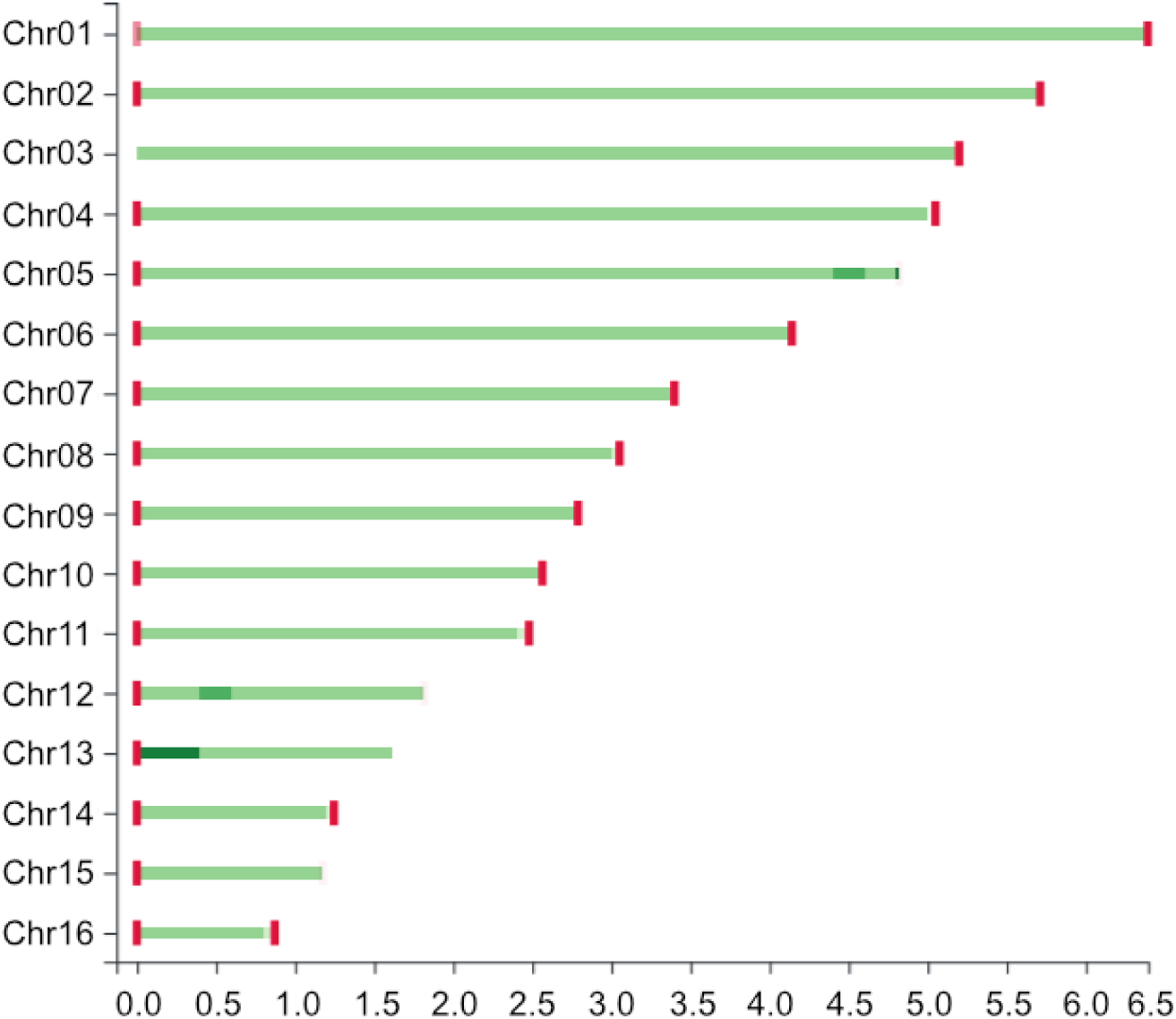
Telomere identification across the 16 assembled chromosomes of *Fusarium oxysporum* MRL8996. Horizontal bars represent the individual chromosomes of the MRL8996 genome. Green bars indicate the sizes of the scaffolds, while variations in colour intensity reflect the read coverage. The read coverage for each scaffold is determined by mapping Nanopore reads onto the assembly. Red vertical lines at the ends of the chromosomes indicate the successful detection of telomeric repeats. A total of 27 out of the 32 expected telomeres were identified.

**Supplementary Image 2.**
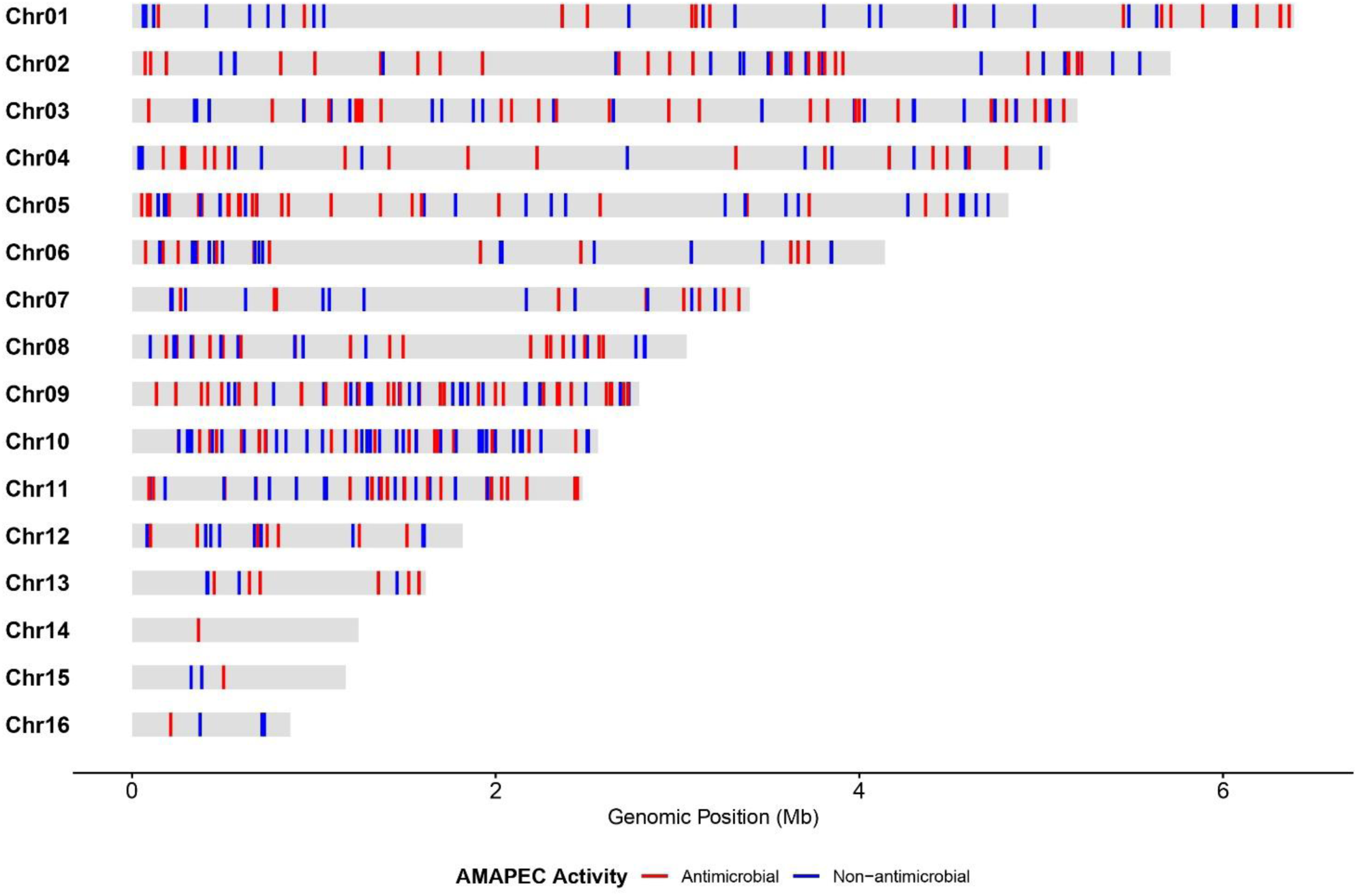
Whole-genome distribution of the MRL8996 predicted effector proteins. Linear genomic map showing the location of the 513 predicted effector genes. Chromosomes are represented by grey bars. Blue vertical lines indicate predicted non-antimicrobial effectors, while red vertical lines indicate predicted antimicrobial effectors.

**Supplementary Image 3.**
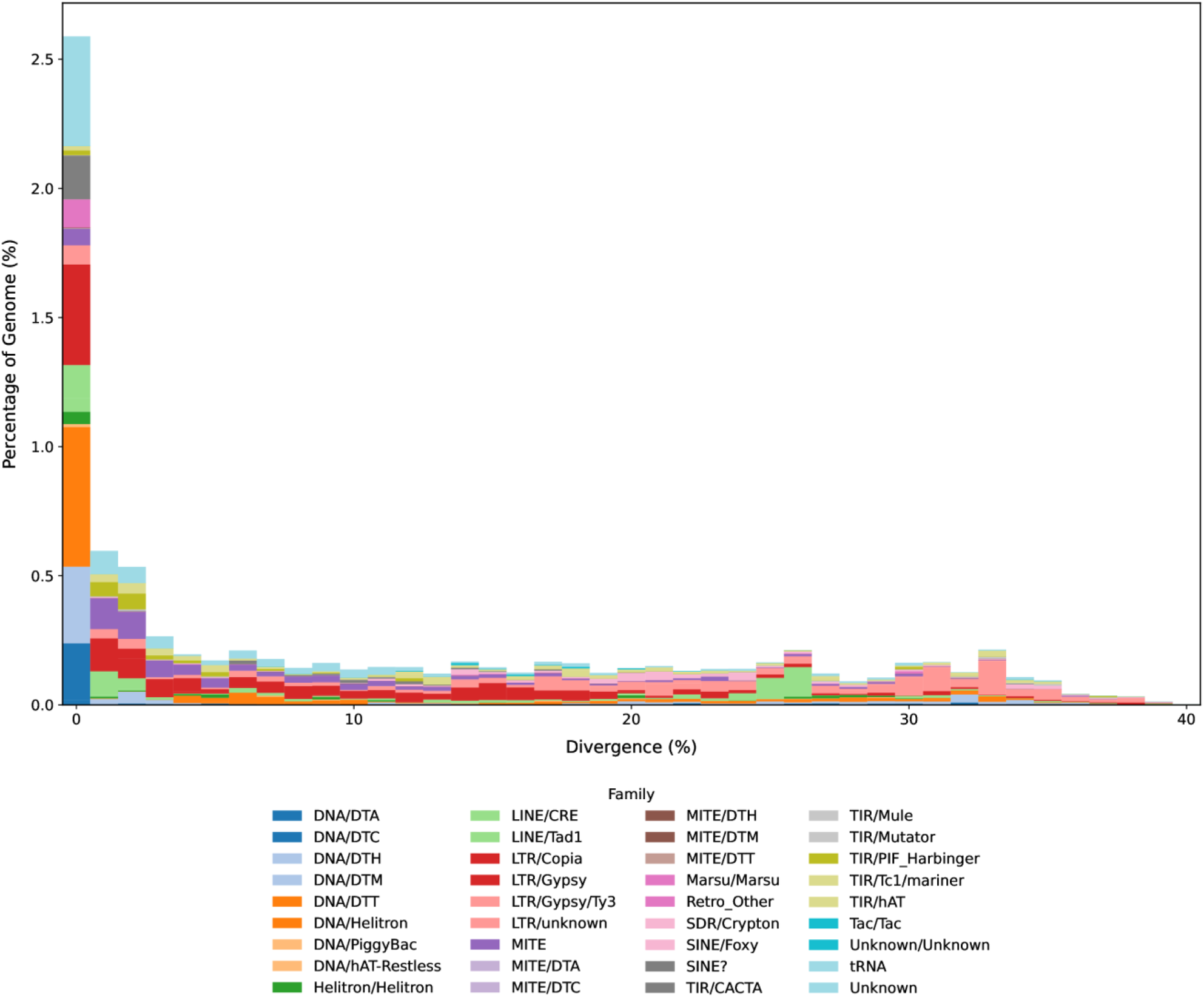
Characterisation of transposable elements across *Fusarium oxysporum* MRL8996 chromosomes. The stacked bar chart shows the distribution of TE sequences according to their Kimura substitution levels (divergence from the consensus sequence). The y-axis shows the percentage of the MRL8996 genome occupied by each TE class, while the x-axis indicates the Kimura divergence percentage, serving as a proxy for time elapsed since TE duplication events (0% representing very recent events).

**Supplementary Image 4.**
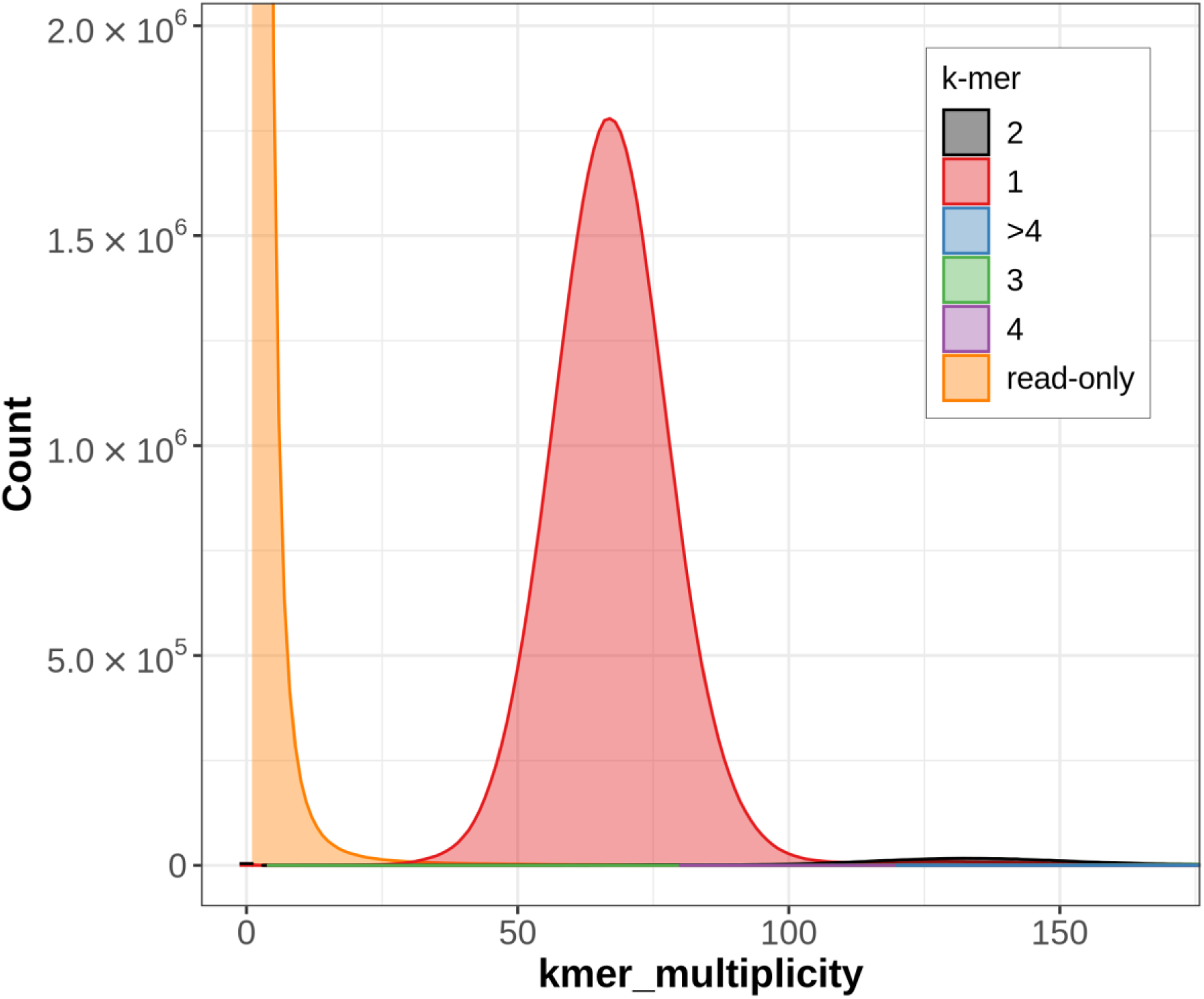
Analysis of k-mer spectra using Merqury to evaluate the completeness and base-level accuracy of the chromosome-level genome assembly of *Fusarium oxysporum* MRL8996. The graph illustrates the k-mer (k=21) multiplicity plot generated by Merqury, which is used to assess the assembly’s completeness and accuracy. The single red peak indicates the presence of single-copy k-mers, suggesting that the genome is highly complete and accurate, with no artefactual duplications. Additionally, the orange “read-only” peak near zero corresponds to sequencing errors that have been effectively excluded from the final sequence.

